# Integrating Developmental Neurotoxicity Prediction with Adverse Outcome Pathway Reasoning

**DOI:** 10.64898/2026.09.28.755098

**Authors:** Yihan Fan, Shuo Xiao, Feng Gao

**Author notes:** Corresponding Author: Feng Gao, PhD, Department of Environmental Health Sciences, Fielding School of Public Health, and Department of Molecular and Medical Pharmacology, David Geffen School of Medicine, University of California, Los Angeles, Los Angeles, CA, USA.

## Abstract

Experimental assessment of developmental neurotoxicity (DNT) is costly, time-consuming, and difficult to scale across large chemical inventories. New approach methodologies (NAMs), including computational approaches based on machine learning, deep learning, and large language models (LLMs), offer new ways to screen and prioritize chemicals. However, these emerging computational approaches have largely focused on predicting toxicity endpoints, with less attention to the mechanisms underlying those predictions. For DNT assessment, understanding this mechanistic basis is equally important. In this study, we integrated LLM inference with the adverse outcome pathway (AOP) framework to connect DNT predictions with plausible mechanisms. We constructed an AOP-informed benchmark linking ChEMBL target-activity measurements for 5,861 compounds to 14 AOPs representing five neurodevelopmental outcomes and evaluated ten LLMs for the prediction of molecular initiating events (MIEs), key events (KEs), key event relationships (KERs), and adverse outcomes (AOs), as well as the generation of mechanistic explanations. Explanation quality was evaluated using both a naive LLM judge and a rubric-based LLM judge. Across the zero-shot models, the highest observed macro-F1 scores were 0.818 for MIEs and 0.463 for AOs, but only 0.056 for KEs and 0.034 for KERs, identifying downstream pathway reconstruction as the principal challenge. Few-shot prompting substantially improved performance, with Gemini 3.8 Flash under four-shot prompting achieving macro-F1 scores of 0.956 for AOs, 0.813 for MIEs, 0.492 for KEs, and 0.457 for KERs. Both LLM-judge configurations indicated improved mechanistic explanation quality with few-shot examples, although strong outcome prediction did not consistently ensure recovery of the recorded pathway. Our results demonstrate the potential of AOP-informed LLMs to support DNT assessment by linking predicted neurodevelopmental outcomes to plausible biological mechanisms and enabling predictions to be evaluated at both the outcome and pathway levels.

## 1. Introduction

Neurodevelopment encompasses the formation and maturation of the nervous system, providing the foundation for cognition, behavior, and motor function ^1^. The developing nervous system is particularly vulnerable to disruption by environmental chemicals during sensitive periods from prenatal development through adolescence ^2^. Such disturbances can have lasting neurological consequences, making developmental neurotoxicity (DNT) an important environmental health concern ^3^. However, conventional DNT studies are costly, time-consuming, and resource-intensive, limiting the number of chemicals that can be evaluated experimentally ^4, 5^. New approach methodologies (NAMs), including in vitro systems and computational models based on machine learning (ML) and deep learning (DL), provide opportunities to expand chemical screening, integrate diverse evidence, and prioritize chemicals for further testing ^6–11^.

Most ML and DL toxicity models are designed primarily to predict an endpoint, such as whether a chemical is likely to produce a particular toxic effect ^6, 7, 9, 11–13^. Although endpoint prediction is useful for screening, it does not establish whether the prediction is supported by a biologically plausible mechanism. This distinction is particularly important for DNT, which can arise through multiple molecular targets and downstream developmental processes. Adverse outcome pathways (AOPs) provide a structured framework for connecting a molecular initiating event (MIE) to downstream key events (KEs) and an adverse outcome (AO) through directed key event relationships (KERs) ^14–16^. By organizing evidence across levels of biological organization, AOPs can support the evaluation of both a predicted DNT outcome and the mechanistic explanation linking chemical activity to that outcome. Applying this framework to individual chemicals requires identifying relevant molecular targets, determining the direction of perturbation, and recovering the biological events through which that perturbation may contribute to a neurodevelopmental effect.

Large language models (LLMs) have recently been applied to molecular property and toxicity prediction ^17–21^. Early studies focused largely on endpoint prediction. For instance, Yang et al. compared general-purpose LLMs with conventional machine learning and deep learning models for several toxicity endpoints and further explored potential cardiotoxic mechanisms using molecular docking ^21^. More recent studies have begun to incorporate biological context and explicit mechanistic explanations. CoTox combined chemical structures with pathway and Gene Ontology information to support stepwise reasoning for organ-specific toxicity prediction ^22^. ToxReason introduced an AOP-based benchmark that evaluates both organ-level toxicity prediction and mechanistic reasoning from MIEs to AOs, demonstrating that strong predictive performance does not necessarily correspond to biologically reliable reasoning ^23^. However, these studies did not systematically evaluate DNT-specific outcomes or the developmental mechanisms connecting molecular perturbations to those outcomes. It therefore remains unclear how accurately LLMs can identify DNT-related MIEs, recover downstream KEs and KERs, and predict neurodevelopmental outcomes within a common pathway-based evaluation framework. It is also unclear whether few-shot examples improve DNT-related predictions and reasoning capabilities^23^.

In this study, we constructed a DNT-focused benchmark by linking chemical target-activity measurements from ChEMBL to AOP-Wiki pathways associated with five neurodevelopmental adverse outcomes ^24, 25^. We evaluated ten LLMs on four connected tasks: MIE inference, generation of MIE–KE–AO mechanistic explanations, and AO prediction. We also compared zero-shot, one-, two-, and four-shot prompting to assess how examples influence prediction and explanation performance. Finally, we assessed mechanistic explanation quality using a naive-LLM judge and a rubric-based LLM judge. This study evaluates the potential of general LLMs to support mechanistically informed DNT assessment by integrating chemical activity, biological pathways, and adverse outcome prediction.

## 2. Methods

### 2.1. Dataset construction

#### 2.1.1. Selection of adverse outcomes, pathways, and molecular initiating events

We selected five DNT-related AOs (i.e., learning and memory impairment, decreased cognitive function, decreased intellectual quotient, impaired locomotor function, and increased risk of microcephaly) and their associated AOPs from AOP-Wiki (Release 2.8) ^25^. We retained pathways with explicit developmental relevance in their titles, short names, or life-stage applicability. Within these pathways, we selected MIEs involving activation or inhibition of defined molecular targets and extracted the corresponding KEs and directed KERs. The selected AO–AOP–MIE associations are summarized in Table 1.

**Table 1.**
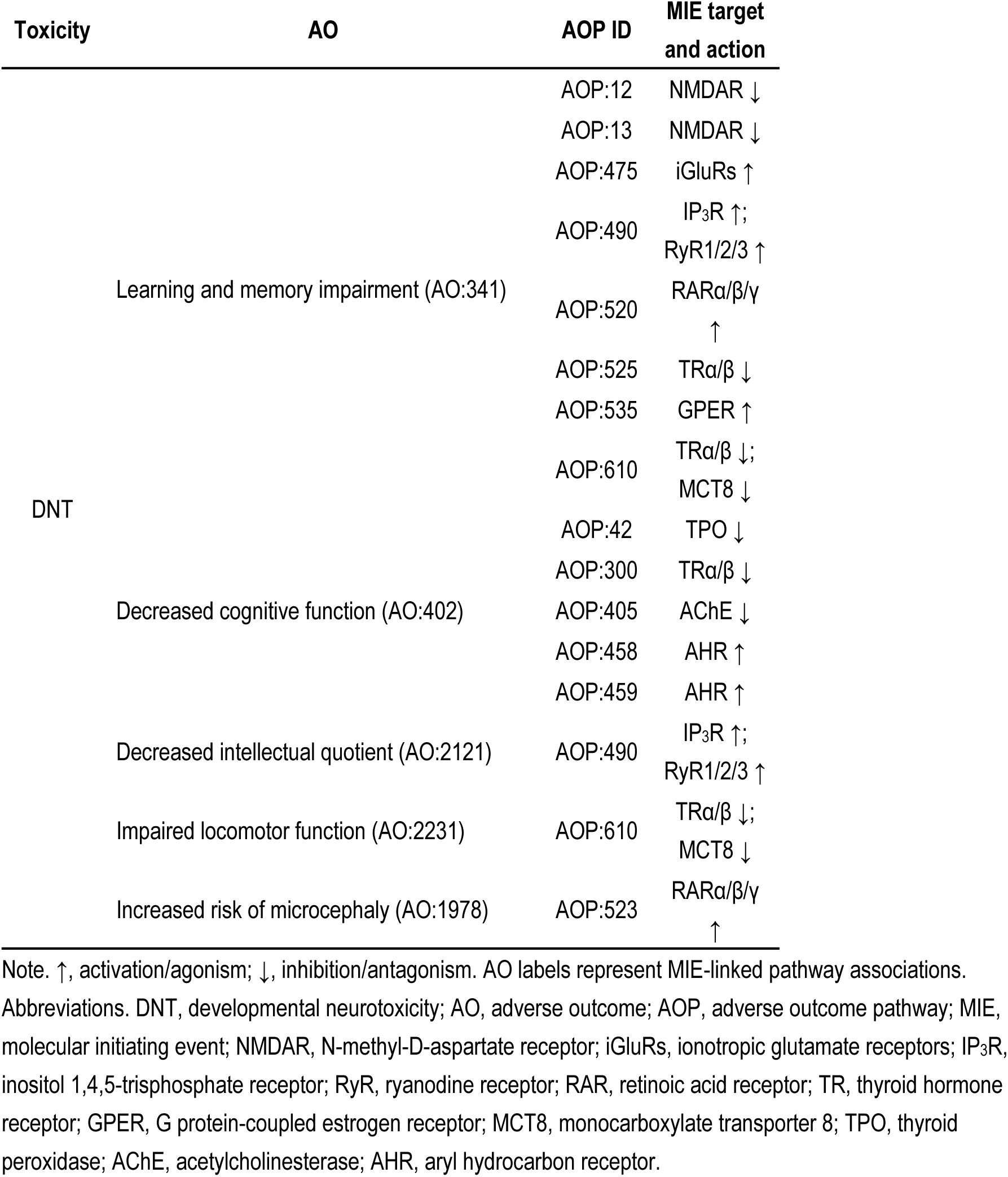
Adverse outcomes, adverse outcome pathways, and molecular initiating events represented in the benchmark.

#### 2.1.2. Derivation of chemical–pathway associations

For each selected MIE, we retrieved activity data for the corresponding human targets from ChEMBL (Release 37) ^24^. Gadaleta et al. used a 10 μM threshold to classify ChEMBL bioactivity records as active or inactive for MIE targets in AOP-based models of organ-specific toxicity, including developing-brain outcomes ^26^. Following this framework and the EC₅₀-for-activation/IC₅₀-for-inhibition implementation used in ToxReason, we classified a compound–target pair as active when at least one qualifying measurement was below 10 μM ^23^. Bioactivity data were aggregated across family members when the MIE was defined at the protein-family level, unless a single isoform was specified.

We combined the selected compounds into a unified dataset, retaining their corresponding targets and directions of action. Compounds were linked to DNT-related AOs through recorded AOP paths connecting their associated MIEs to those outcomes. All qualifying MIE–AOP–AO associations were retained, with a total of 7,837 compound-specific associations across 20 distinct combinations.

#### 2.1.3. Dataset preparation

Chemical structures were standardized using RDKit and represented as canonical isomeric SMILES, with duplicate compounds merged ^27^. The resulting dataset contained 5,861 unique compounds.

The full dataset comprised six mutually exclusive AO-label combinations: 253 compounds annotated only for learning and memory impairment (AO:341), 4,931 only for decreased cognitive function (AO:402), 221 for AO:1978 and AO:341, 63 for AO:2121 and AO:341, one for AO:2231 and AO:341, and 392 for AO:2231, AO:341, and AO:402. To prevent the AO:402-only group from dominating the test set, we randomly selected 50 compounds without replacement from each of the five combinations containing at least 50 compounds and included the sole compound in the remaining group. This yielded a 251-compound test set and the rest 5,610-compound set was used to select few-shot examples and to train the random-forest baseline. The same test set was used throughout the study.

#### 2.1.4. Construction of chemical reference contexts

We constructed a chemical reference library from ChEMBL measurements for the selected MIE targets. For MIEs defined by activation, compounds were annotated as activating or non-activating; for MIEs defined by inhibition, they were annotated as inhibiting or non-inhibiting, using the 10 μM activity cutoff. To retrieve structurally similar chemicals, we represented molecular structures as 1,024-bit Morgan fingerprints with a radius of 2 using RDKit ^28^. Similarity between query and reference compounds was calculated using the Dice coefficient. For each query compound, we excluded the compound itself and records representing the same chemical identity before ranking the references. We selected the four most similar compounds from each of the activation and inhibition groups. Their SMILES, similarity scores, MIE targets, directions of action, and corresponding activity annotations were assembled into a chemical reference context. The query’s reference AO annotations and AOP paths were withheld from model inputs.

### 2.2. Models and experimental settings

We evaluated the performance of ten LLMs under a common zero-shot setting using the test set described in Section 2.1.3. The models included GPT-6 Sol ^29^, GPT-5.4 ^30^, GPT-5.1 ^31^, and GPT-4o ^32^, Claude Sonnet 5 ^33^, Claude Haiku 4.5 ^34^, Gemini 3.8 Flash ^35^, Gemini 3.5 Flash ^36^, Qwen3.7-Flash ^37^, and Qwen3.5-Flash ^38^. We used the same task instructions and structured response format for all models and applied identical output processing and scoring procedures. Temperature was set to 0 for all models except Claude Sonnet 5, which used default sampling with adaptive thinking at low effort. Each experimental condition was run three times on the same test set.

We included random forest classifiers as a structure-based baseline for AO prediction ^39^. Compounds were represented by 1,024-bit Morgan fingerprints with a radius of 2. Tree depth and minimum leaf size were selected by maximizing average precision from five-fold cross-validation using the 5610-compound set.

### 2.3. LLM Inference

We formulated four linked tasks: MIE prediction, KE prediction, explanation of MIE– KE–AO mechanisms, and AO prediction. We used two types of prompts: the system prompt and the compound-specific inference prompt. The system prompt defined these tasks for the five selected DNT-related AOs and specified the required response format. The compound-specific inference prompt provided the query SMILES and its chemical reference context (i.e., the retrieved structurally similar compounds, their similarity scores, and their MIE-related activity annotations). LLMs completed all four tasks in a single structured response containing predicted target-direction pairs, biological events, directed relationships, mechanistic explanations, and AO labels. The system prompt and compound-specific inference prompt used for DNT inference are provided in Table 2.

**Table 2.**
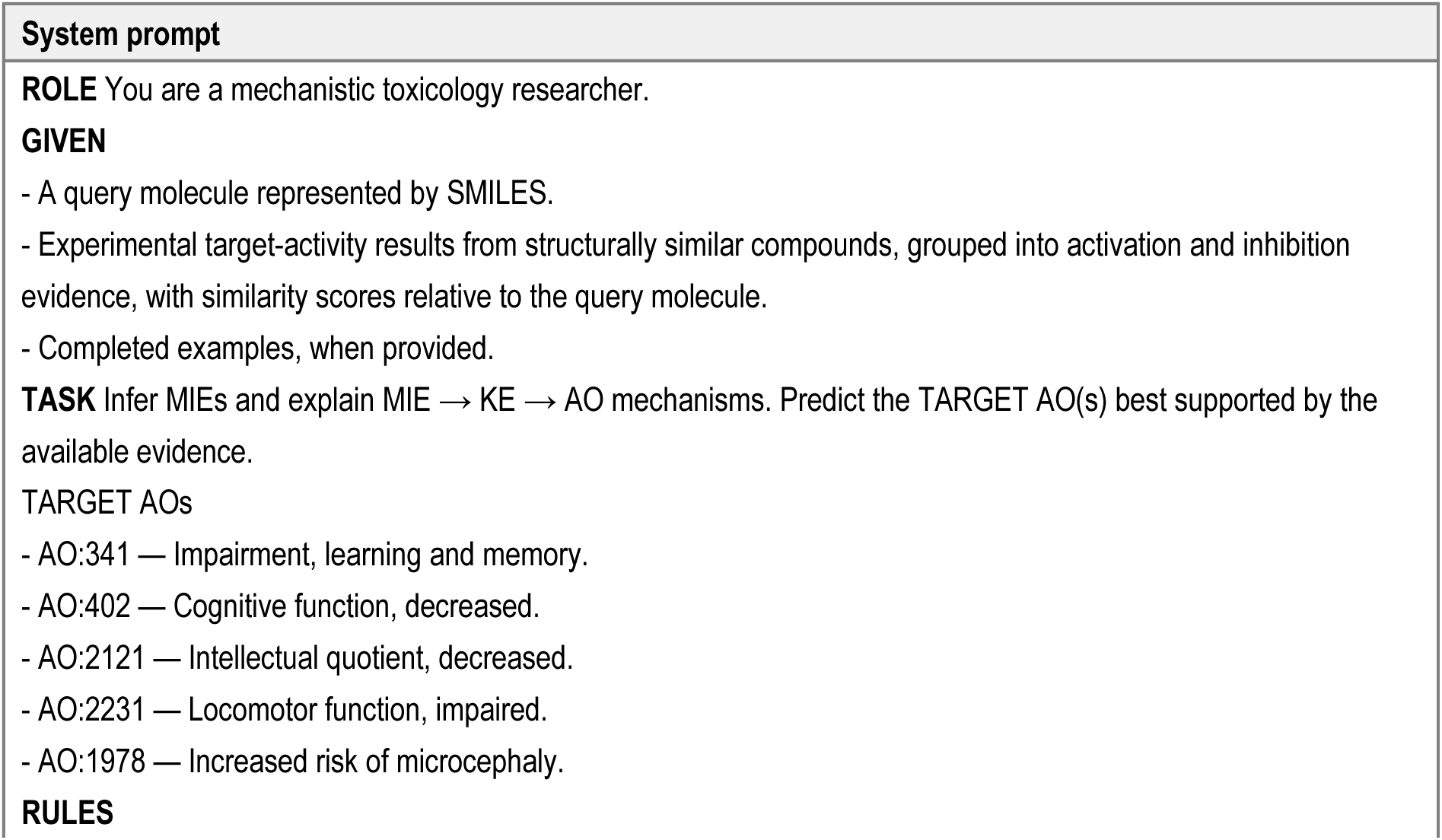
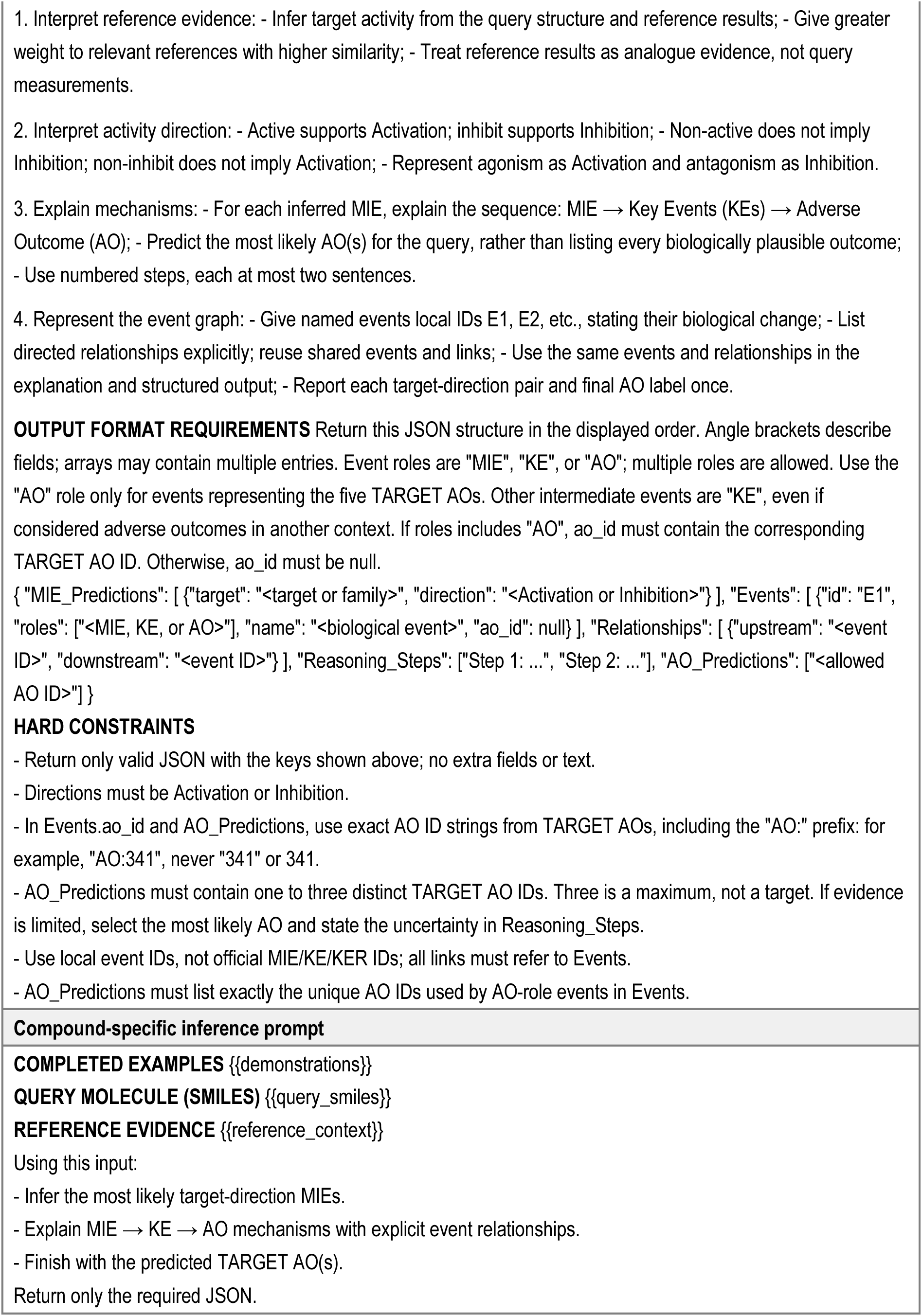
Inference prompts.

Under zero-shot conditions, the prompt contained only the query and its chemical reference context, without any examples. To assess whether providing examples improves DNT-related prediction and mechanistic reasoning, we additionally evaluated few-shot prompting ^40^. We provided GPT-5.1 and Gemini 3.8 Flash with one, two, or four examples randomly selected from the remaining chemical set. Each example paired the compound’s SMILES and reference context with a completed response covering all four tasks. Example MIE and AO labels were obtained from the reference annotations, and the corresponding KEs and directed KERs were assembled from recorded AOP paths. The accompanying mechanistic explanations were generated using GPT-6 Astra ^41^. The examples were inserted before the query in the compound-specific inference prompt. Their content and order were fixed across models and repeated runs, with the one- and two-shot conditions using the first one or two examples from the four-example set.

### 2.4. Evaluation

#### 2.4.1. AO, MIE, KE, and KER prediction

We compared predicted AO, MIE, KE, and KER annotations with the reference annotations for each compound. AOs were matched by identifier. Each predicted MIE was represented as a target–direction pair, with direction specified as activation or inhibition. Predicted target names, abbreviations, and gene symbols were mapped to the corresponding target or family, and an MIE was counted as a match only when both the target and direction agreed with the reference annotation. KEs were matched using official event names or predefined patterns describing the biological entity and direction of change. KER matching required the same upstream and downstream events in the correct order.

We calculated F1 separately for each reference label (i.e., AOs, MIEs, KE, KER) across test compounds. Macro-F1 was then calculated as the unweighted average of the label-specific F1 scores within each category (e.g., the mean F1 across the five AOs for AO macro-F1). Predictions of labels not represented in the predefined reference sets were recorded separately and did not contribute to macro-F1.

#### 2.4.2. Naive LLM-based judge for mechanistic reasoning evaluation

Following Park et al., we evaluated the mechanistic reasoning quality of LLM-generated explanations across multiple dimensions: hallucination avoidance, causal coherence, biological fidelity, and a newly added informativeness dimension ^23^. Hallucination avoidance assessed whether explanations avoided fabricated information, misrepresented evidence, or conclusions exceeding the available support. Causal coherence assessed whether MIEs, KEs, and AOs formed a logically justified sequence without key gaps, reversed relationships, or contradictions. Biological fidelity assessed the accuracy of biological terminology, target functions, directions of action, and mechanistic relationships in the relevant developmental context. Informativeness assessed whether explanations provided specific and relevant mechanistic details that helped explain the predicted outcomes. We used GPT-5.4 mini ^42^ as the LLM judge to evaluate mechanistic explanations and our prompts instructed the judge to assign a score between 0 and 10 for each dimension and provide a brief justification, with higher scores indicating better explanation quality. The judge was instructed to distinguish identifiable errors from claims that could not be verified using the provided information. An overall score was calculated as the arithmetic mean of the four-dimension scores. The system and input prompts used for explanation evaluation are provided in Table 3.

**Table 3.**
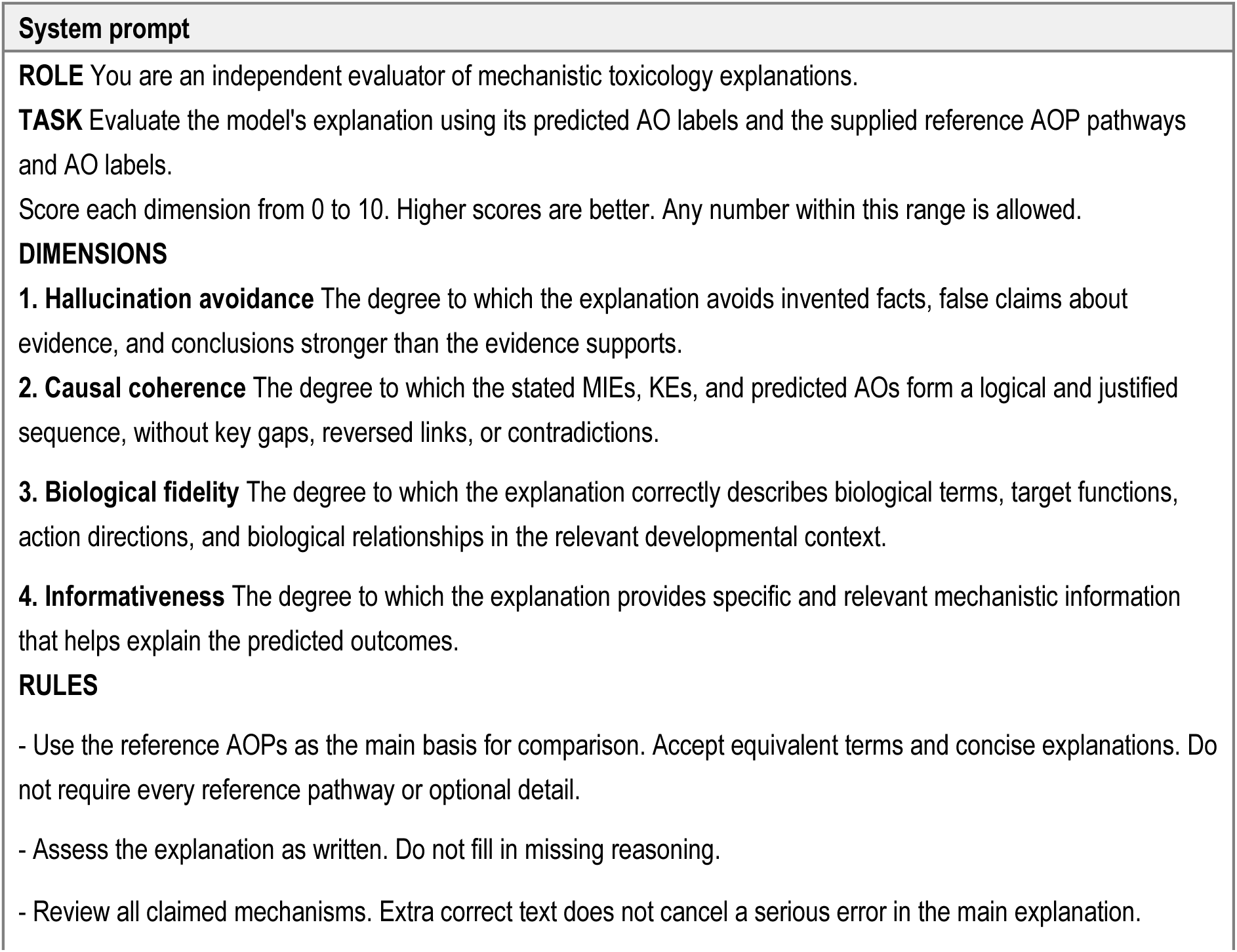
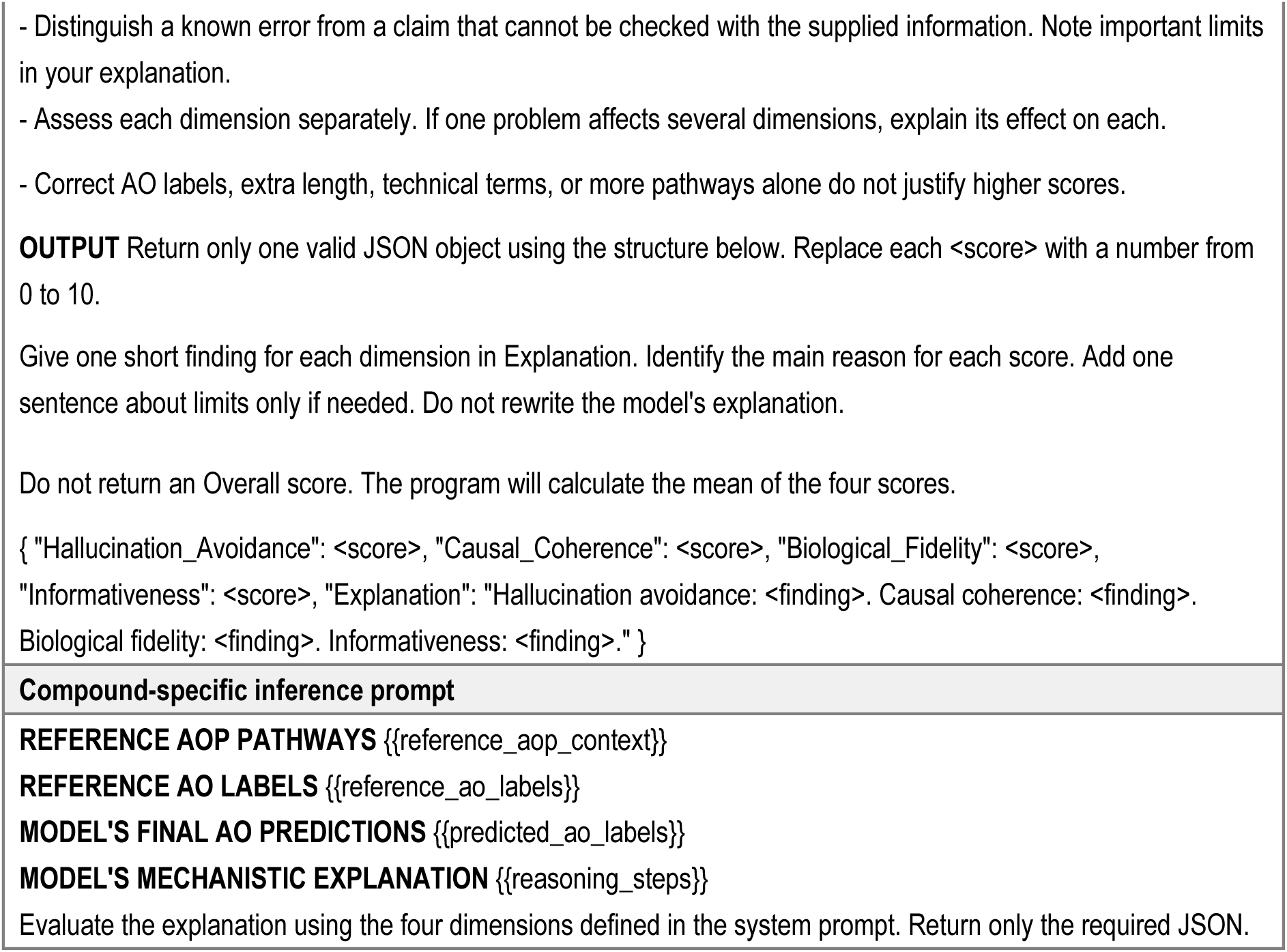
Naive LLM judge prompts.

#### 2.4.3. Rubric-based LLM judge for mechanistic reasoning evaluation

We retained the same LLM judge model and four evaluation dimensions and added explicit dimension-specific scoring criteria to the system prompt. The rubric distinguished errors affecting the central mechanism from those confined to secondary details. It also specified criteria for the completeness of causal connections and the relevance and specificity of mechanistic information. Compound-specific inference prompt, requirement for score justifications, and calculation of the overall score were unchanged. The rubric-guided system and input prompts are provided in Table 4. Figure 1 summarizes the study workflow.

**Figure 1.**
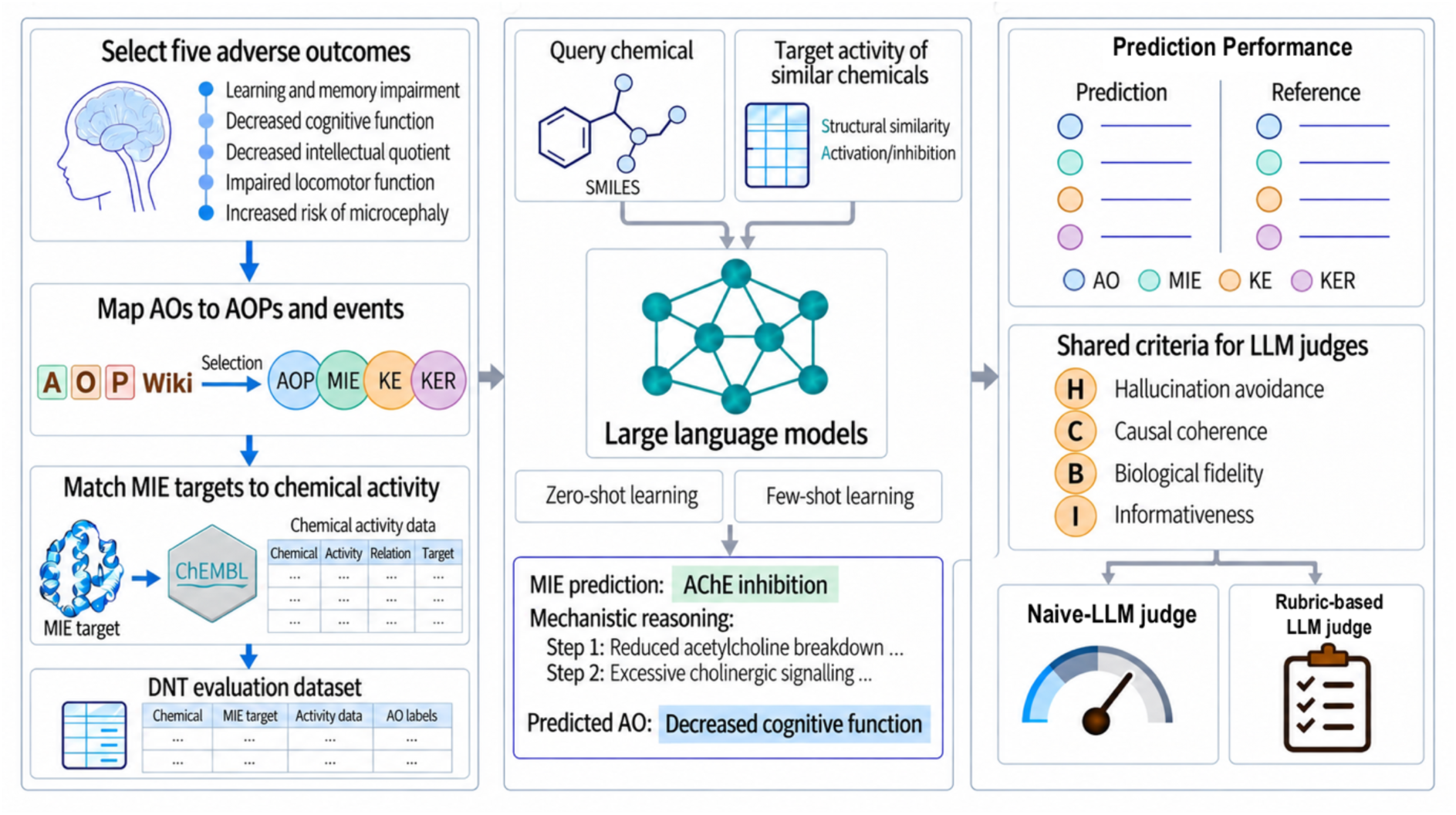
Study workflow. (Left) Construction of the DNT evaluation dataset using AOP-Wiki and ChEMBL. (Middle) LLM inference under zero-shot and few-shot learning settings. (Right) Evaluation through reference matching, the naive-LLM judge and rubric-based LLM judge. **Abbreviations.** AO, adverse outcome; MIE, molecular initiating event; KE, key event; KER, key event relationship

**Table 4.**
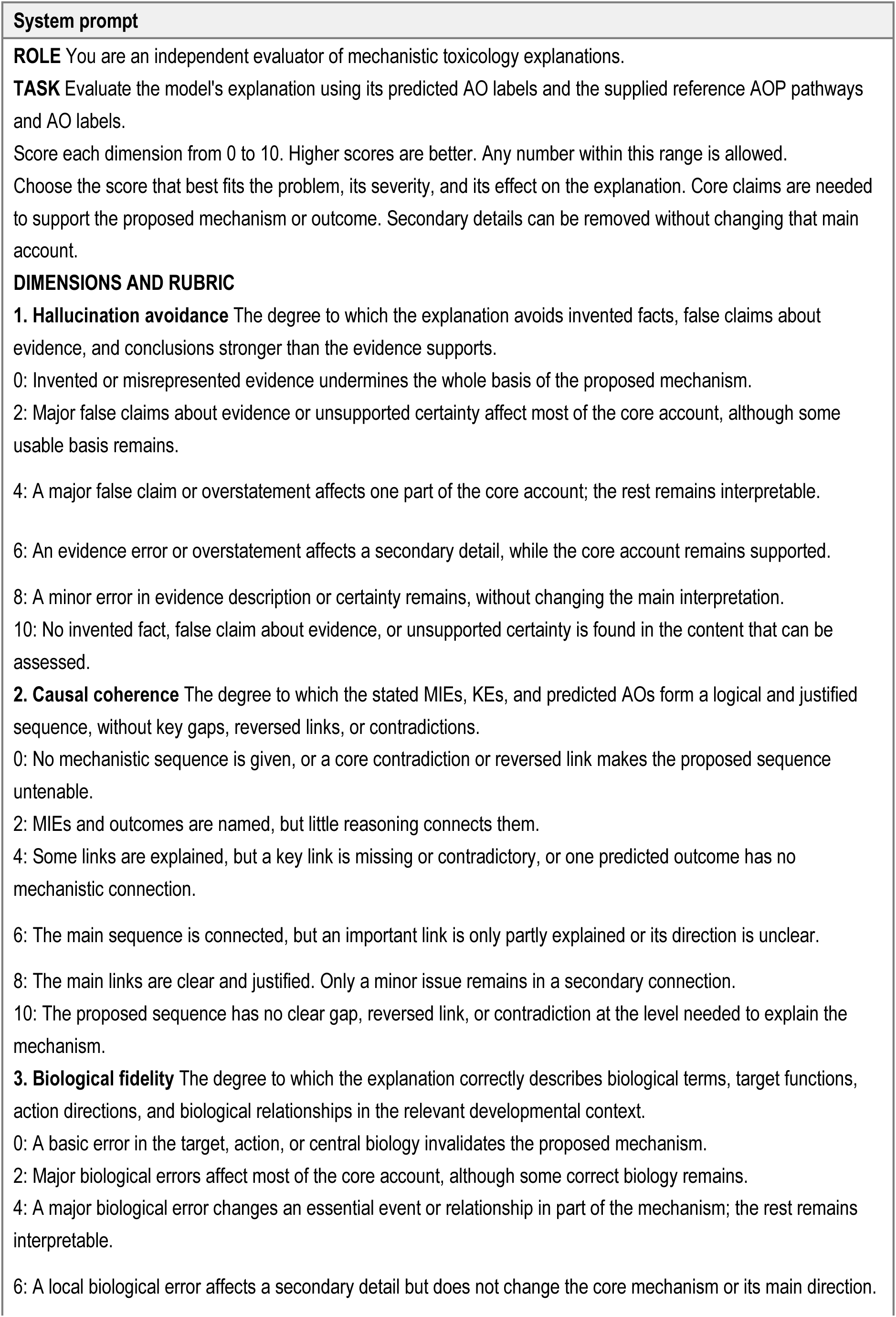
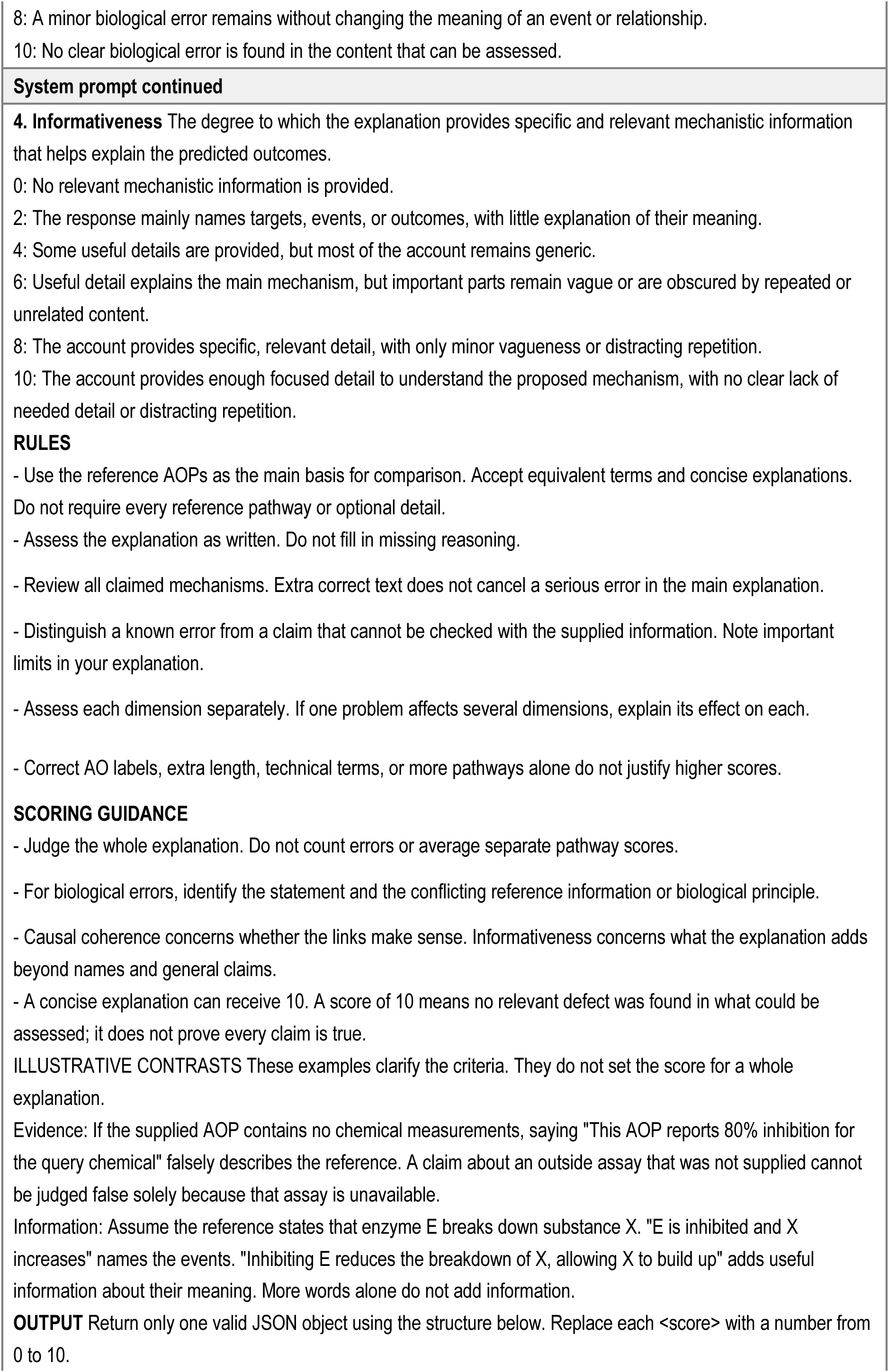
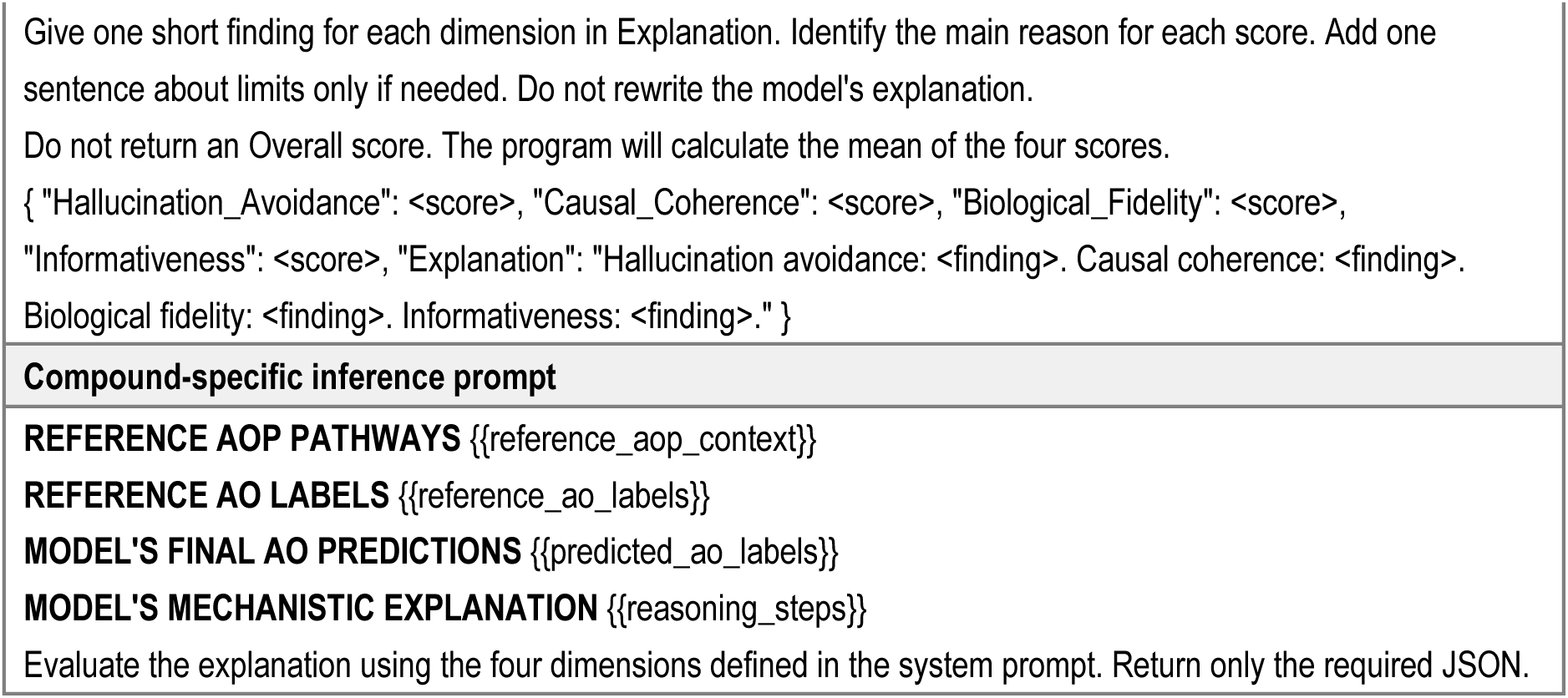
Rubric-based LLM judge prompts.

### 2.5. Statistical analysis

Performance was summarized as the mean and sample standard deviation across three experimental repeats. LLM-judge scores were first averaged across compounds within each repeat. For comparisons between the two judge approaches, scores were averaged across repeats for each compound. Spearman rank correlations were then calculated across compounds to assess ranking consistency, separately for each model condition, scoring dimension, and overall score. Paired score differences were calculated as rubric-based LLM judge minus naive-LLM judge.

## 3. Results and Discussion

### 3.1. DNT-related pathway coverage and dataset characteristics

The dataset comprised 5,861 compounds linked to five DNT-related AOs through 14 AOPs (Table 1). Several AOs shared MIEs. For example, retinoic acid receptor activation occurred in pathways associated with both learning and memory impairment (AO:341) and increased risk of microcephaly (AO:1978). These overlapping pathway associations produced a multilabel prediction setting, with 677 compounds (11.6%) carrying more than one AO annotation. The test-set reference annotations comprised five AO labels, 10 MIE target–direction labels, 49 KEs, and 99 directed KERs.

### 3.2. Zero-shot performance in DNT-related AO prediction and mechanistic explanation

Under zero-shot conditions, the ten LLMs performed better on MIE prediction than AOs or downstream pathway components (Table 5). Macro-F1 ranged from 0.557 to 0.818 for MIEs and from 0.261 to 0.463 for AOs, with GPT-6 Sol achieving the highest MIE performance and Claude Sonnet 5 and Gemini 3.8 Flash sharing the highest AO performance, respectively. Prediction of the downstream pathway structure was substantially weaker, with KE and KER macro-F1 ranging from 0.002 to 0.056 and from 0 to 0.034, respectively. Thus, relatively strong prediction at the molecular initiation stage coexisted with limited recovery of the downstream events and relationships. The larger KE and KER reference pool and the stricter requirement for correctly ordered event pairs may partly contribute to this pattern.

**Table 5.** Prediction and mechanistic reference-matching performance across three experimental repeats.

| Model | AO | MIE | KE | KER |
| --- | --- | --- | --- | --- |
|  | Macro-F1 | Macro-F1 | Macro-F1 | Macro-F1 |
| GPT-6 Sol | 0.350(0.005) | 0.818(0.000) | 0.025(0.002) | 0.008(0.000) |
| GPT-5.4 | 0.402(0.016) | 0.743(0.001) | 0.039(0.003) | 0.012(0.002) |
| GPT-5.1 | 0.424(0.027) | 0.614(0.004) | 0.050(0.001) | 0.018(0.001) |
| GPT-4o | 0.261(0.010) | 0.557(0.019) | 0.002(0.002) | 0.000(0.000) |
| Claude Sonnet 5 | 0.463(0.012) | 0.791(0.007) | 0.034(0.002) | 0.014(0.001) |
| Claude Haiku 4.5 | 0.373(0.008) | 0.778(0.001) | 0.050(0.001) | 0.034(0.003) |
| Gemini 3.8 Flash | 0.463(0.007) | 0.804(0.001) | 0.056(0.005) | 0.022(0.000) |
| Gemini 3.5 Flash | 0.449(0.009) | 0.788(0.021) | 0.050(0.004) | 0.017(0.001) |
| Qwen 3.7 Flash | 0.269(0.009) | 0.792(0.004) | 0.039(0.002) | 0.014(0.001) |
| Qwen 3.5 Flash | 0.336(0.004) | 0.721(0.018) | 0.021(0.001) | 0.010(0.000) |
| GPT-5.1-1shot | 0.478(0.009) | 0.739(0.052) | 0.066(0.007) | 0.026(0.002) |
| GPT-5.1-2shot | 0.513(0.005) | 0.714(0.058) | 0.138(0.010) | 0.097(0.004) |
| GPT-5.1-4shot | 0.792(0.012) | 0.774(0.059) | 0.359(0.022) | 0.262(0.018) |
| Gemini 3.8 Flash-1shot | 0.488(0.002) | 0.809(0.001) | 0.075(0.001) | 0.054(0.002) |
| Gemini 3.8 Flash-2shot | 0.531(0.003) | 0.808(0.003) | 0.165(0.002) | 0.149(0.001) |
| Gemini 3.8 Flash-4shot | 0.956(0.004) | 0.813(0.005) | 0.492(0.005) | 0.457(0.002) |
| ECFP4-RF | 0.915(0.002) | - | - | - |

Note. Values are mean (SD) across three repeats (251 test compounds each). The ECFP4-RF comparison was restricted to AO prediction. Dashes indicate outcomes not included in the baseline comparison. Abbreviations. AO, adverse outcome; MIE, molecular initiating event; KE, key event; KER, key event relationship; ECFP4-RF, extended-connectivity fingerprint/random forest baseline.

Alternative descriptions that are not recognized by these rules may therefore remain unmatched.

The prediction performance difference between MIE and downstream pathway reflects the information available to the models and the complexity of the corresponding inference tasks. Although no completed examples were provided in the zero-shot setting, each query included structurally similar reference compounds with target, activity-direction, and activity-classification information. The relatively high MIE prediction performance therefore indicates that the models could use this chemical reference context to identify plausible proximal target perturbations. By contrast, the reference context did not provide the downstream events linking those perturbations to DNT related AOs. Recovering KEs and KERs required the models to select among overlapping and branching AOPs, identify developmentally relevant intermediate events, and preserve the direction and order of their relationships. This downstream reasoning step represented the principal bottleneck under zero-shot prompting, requiring both pathway selection and event-level alignment across a substantially larger reference space.

AO prediction also varied substantially across developmental outcomes. Gemini 3.8 Flash and Claude Sonnet 5 shared the highest AO macro-F1 of 0.463. However, performance within a given model was not uniform across outcomes. For example, Gemini 3.8 Flash achieved an F1 of 0.983 for increased risk of microcephaly, but 0 for decreased intellectual quotient and 0.013 for impaired locomotor function. High performance for one developmental AO therefore did not generalize consistently across the DNT outcomes represented in the benchmark. MIE prediction performance also did not consistently translate into AO prediction performance. GPT-6 Sol achieved the highest MIE macro-F1 of 0.818 but an AO macro-F1 of 0.350, while Qwen 3.7 Flash achieved an MIE macro-F1 of 0.792 but an AO macro-F1 of 0.269. Several MIEs occur in multiple AOPs, and some are associated with more than one developmental outcome. Identifying a plausible target-direction pair is therefore insufficient to determine the relevant AO without additional pathway-level information.

Explanation assessment revealed additional differences among models that were not reflected by AO agreement alone. GPT-6 Sol received the highest overall explanation scores under both the Naive-LLM judge and Rubric-based LLM judge, with mean scores of 5.973 and 5.999, respectively (Tables 6 and 7), despite an AO macro-F1 of 0.350. Explanation profiles also differed across scoring dimensions. For example, under the Naive-LLM judge, GPT-5.1 received an informativeness score of 6.031, compared with 3.080 for hallucination avoidance and 3.843 for biological fidelity (Table 6). Thus, greater assessed informativeness did not necessarily coincide with similarly favorable assessments of hallucination avoidance or biological fidelity. A related dissociation between predictive performance and mechanistic explanation quality was reported by ToxReason for liver, heart, and kidney toxicity ^23^.

**Table 6.**
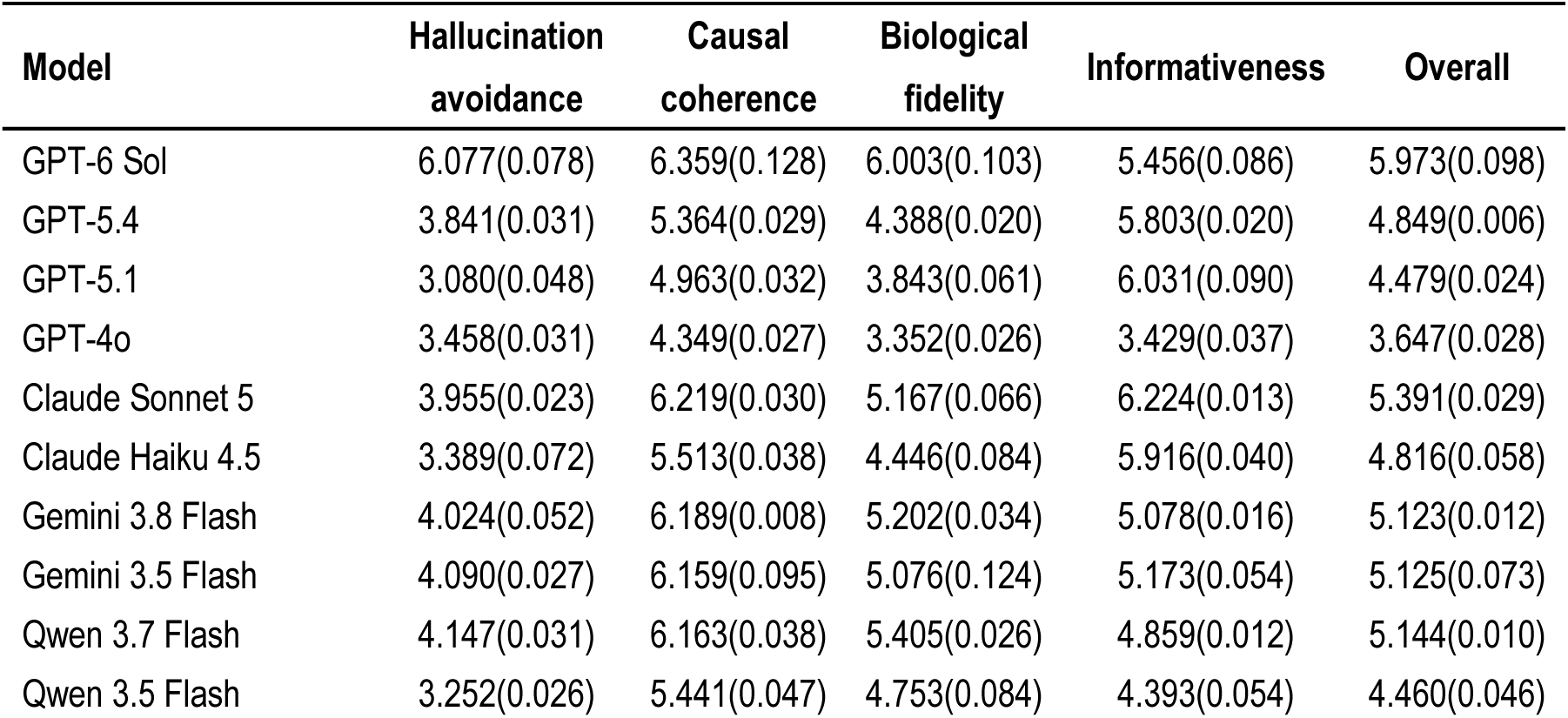
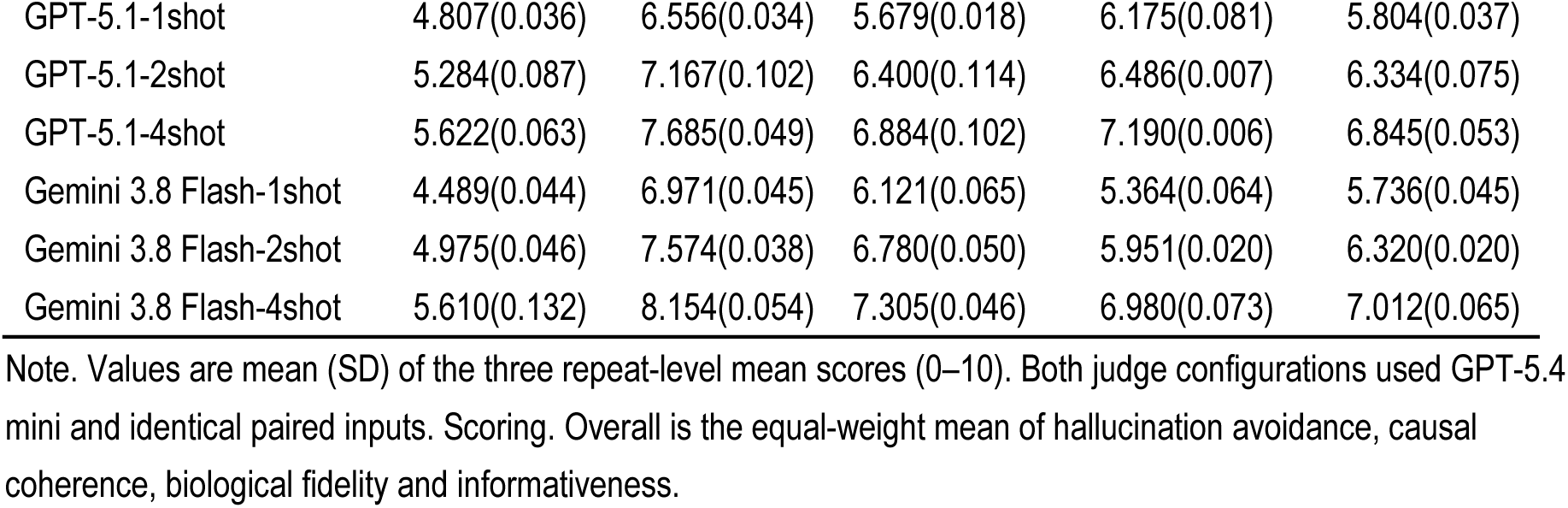
Evaluation of mechanistic explanations using the Naive-LLM judge.

**Table 7.** Evaluation of mechanistic explanations using the Rubric-based LLM judge.

| Model | Hallucination avoidance | Causal coherence | Biological fidelity | Informativeness | Overall |
| --- | --- | --- | --- | --- | --- |
| GPT-6 Sol | 5.198 (0.144) | 6.660 (0.014) | 6.005 (0.085) | 6.133 (0.028) | 5.999 (0.064) |
| GPT-5.4 | 3.236 (0.023) | 5.558 (0.028) | 4.389 (0.054) | 6.827 (0.022) | 5.003 (0.019) |
| GPT-5.1 | 2.675 (0.034) | 5.351 (0.045) | 3.857 (0.086) | 7.060 (0.083) | 4.735 (0.016) |
| GPT-4o | 3.224 (0.089) | 4.663 (0.019) | 3.487 (0.034) | 4.177 (0.036) | 3.888 (0.018) |
| Claude Sonnet 5 | 3.262 (0.062) | 6.434 (0.007) | 5.228 (0.027) | 6.965 (0.010) | 5.472 (0.019) |
| Claude Haiku 4.5 | 2.582 (0.049) | 5.741 (0.017) | 4.331 (0.016) | 6.871 (0.018) | 4.881 (0.021) |
| Gemini 3.8 Flash | 3.041 (0.043) | 6.372 (0.038) | 5.325 (0.042) | 6.008 (0.011) | 5.187 (0.024) |
| Gemini 3.5 Flash | 3.084 (0.069) | 6.242 (0.056) | 5.092 (0.062) | 6.109 (0.050) | 5.131 (0.034) |
| Qwen 3.7 Flash | 3.222 (0.047) | 6.499 (0.072) | 5.749 (0.060) | 5.704 (0.052) | 5.293 (0.054) |
| Qwen 3.5 Flash | 2.583 (0.086) | 5.841 (0.049) | 4.911 (0.022) | 5.390 (0.046) | 4.681 (0.047) |
| GPT-5.1-1shot | 3.955 (0.128) | 6.907 (0.034) | 5.886 (0.047) | 6.989 (0.032) | 5.934 (0.039) |
| GPT-5.1-2shot | 4.202 (0.042) | 7.349 (0.142) | 6.458 (0.165) | 7.145 (0.015) | 6.289 (0.084) |
| GPT-5.1-4shot | 4.301 (0.116) | 7.742 (0.081) | 6.873 (0.127) | 7.639 (0.016) | 6.639 (0.073) |
| Gemini 3.8 Flash-1shot | 3.252 (0.079) | 7.337 (0.023) | 6.501 (0.046) | 6.232 (0.030) | 5.831 (0.020) |
| Gemini 3.8 Flash-2shot | 3.610 (0.066) | 7.867 (0.013) | 7.113 (0.046) | 6.681 (0.021) | 6.318 (0.026) |
| Gemini 3.8 Flash-4shot | 4.210 (0.185) | 8.351 (0.050) | 7.611 (0.051) | 7.473 (0.012) | 6.911 (0.055) |

Note. Values are mean (SD) of the three repeat-level mean scores (0–10). Both judge configurations used GPT-5.4 mini and identical paired inputs. Scoring. Overall is the equal-weight mean of hallucination avoidance, causal coherence, biological fidelity and informativeness.

**Table 8.**
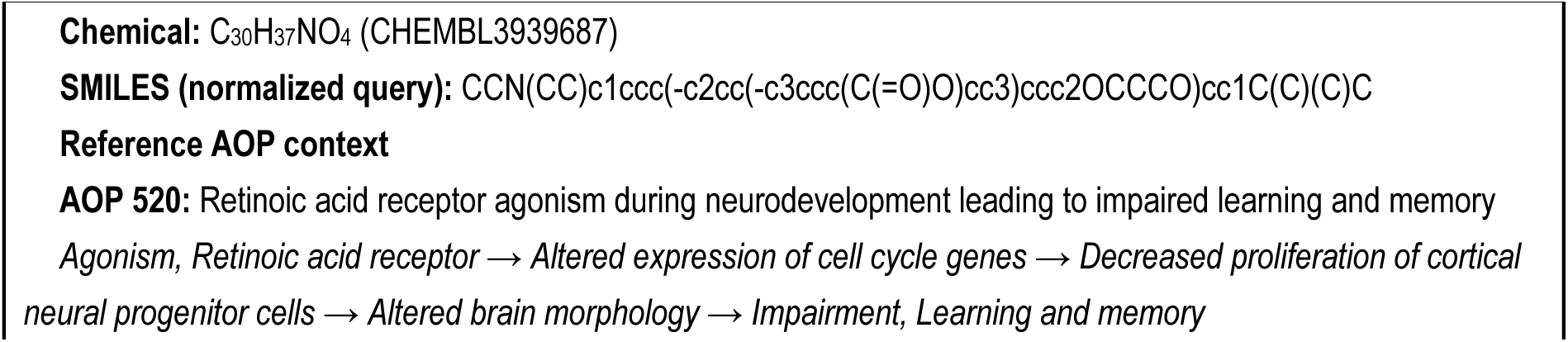
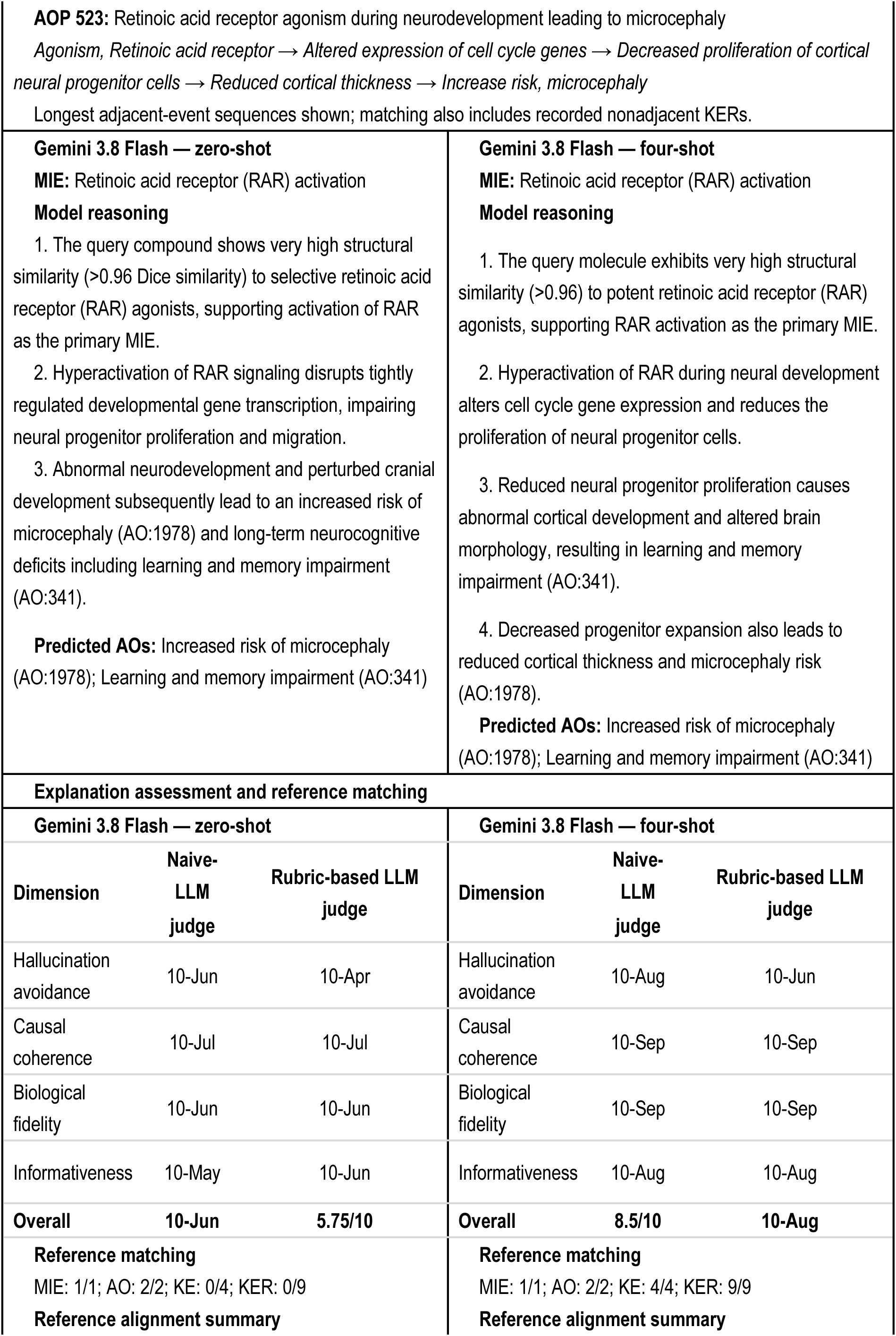
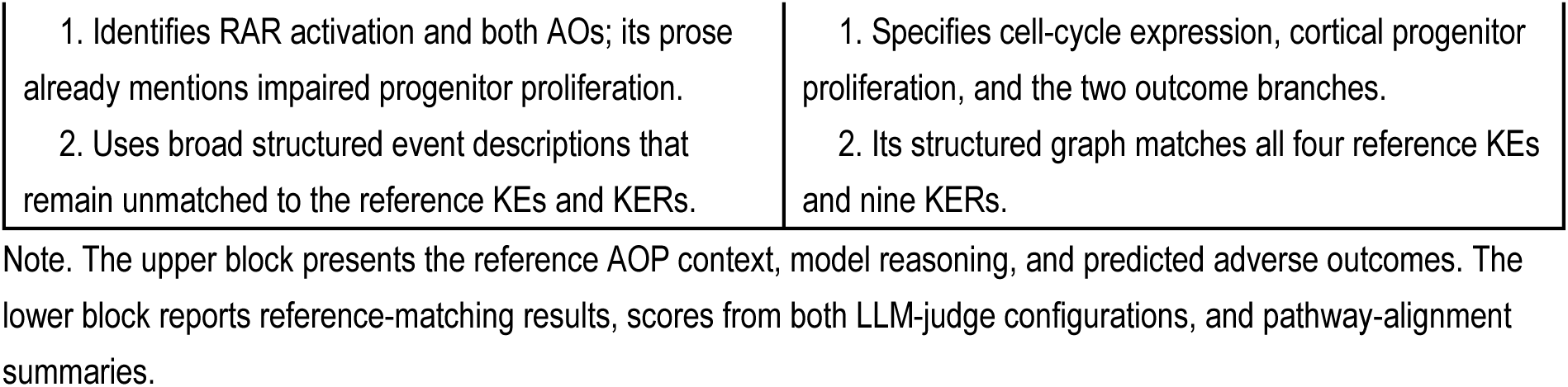
Case study comparing zero-shot and four-shot responses generated by Gemini 3.8 Flash for CHEMBL3939687.

**Table 9.**
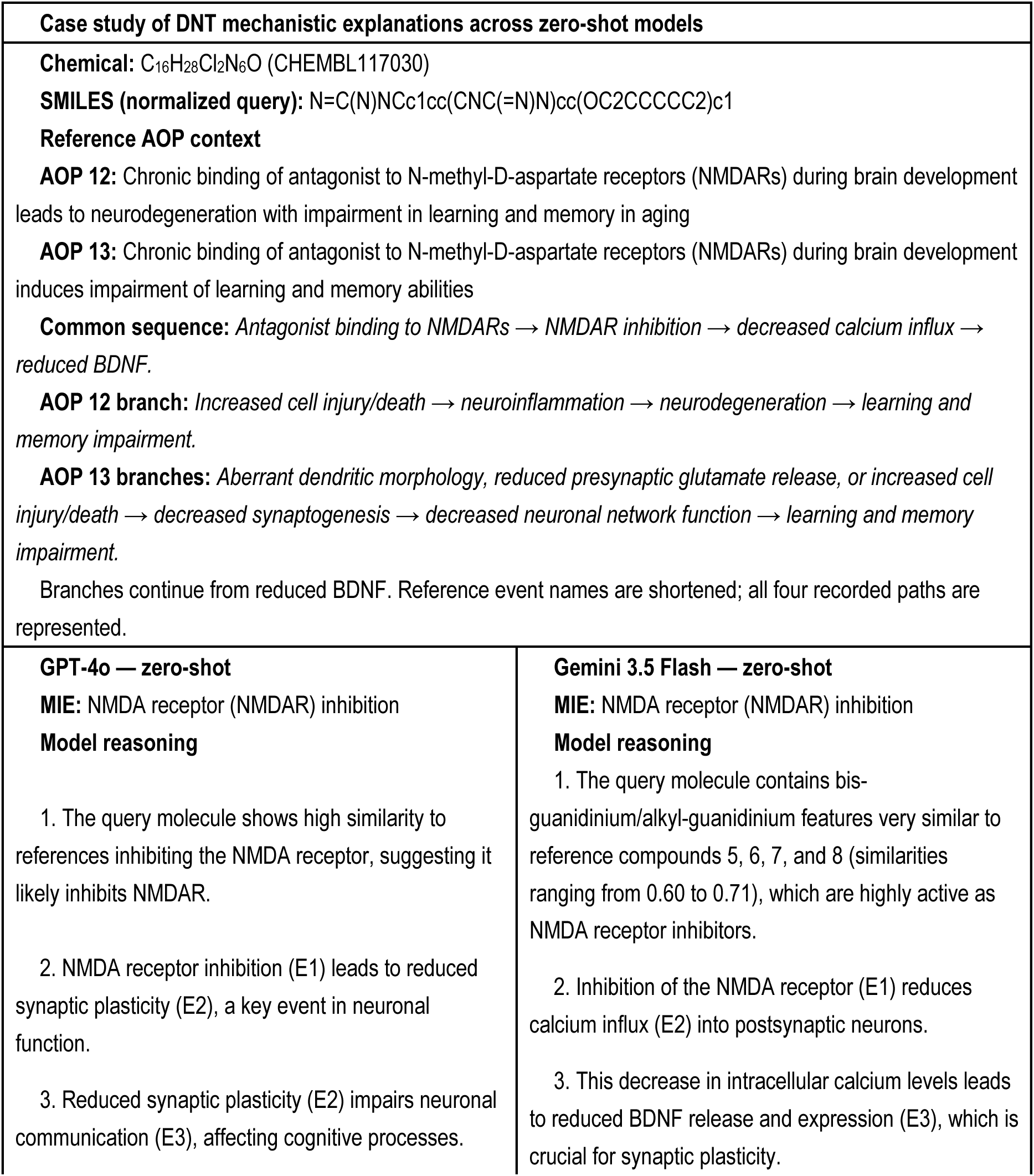
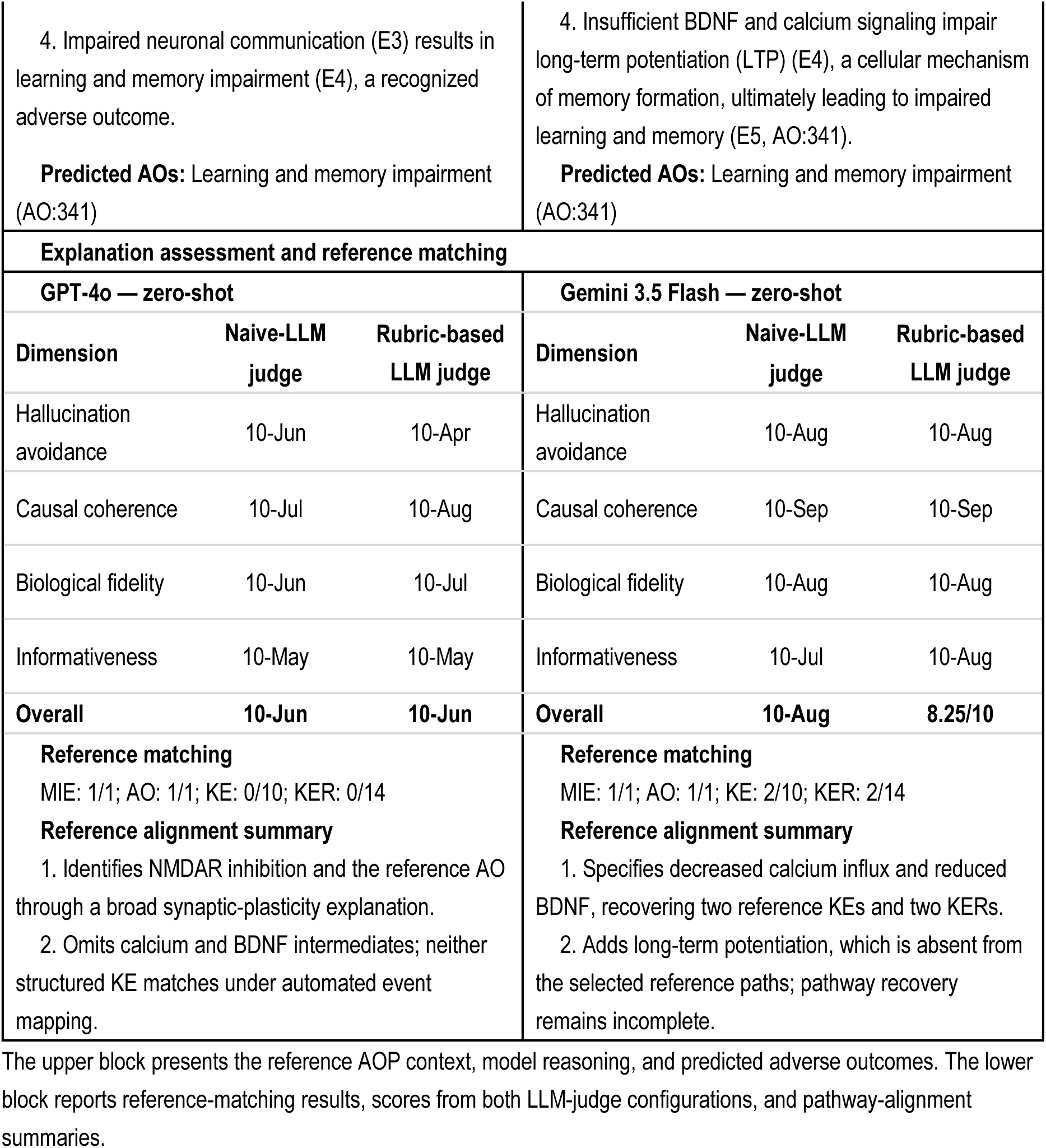
Case study comparing zero-shot responses generated by GPT-4o and Gemini 3.5 Flash for CHEMBL117030.

| Case study of DNT mechanistic explanations across zero-shot models |  |
| --- | --- |
| **Chemical:** C<sub>16</sub>H<sub>28</sub>Cl<sub>2</sub>N<sub>6</sub>O (ChEMBL117030)<br>**SMILES (normalized query):** N=C(N)NCc1cc(CNC(=N)N)cc(OC2CCCCC2)c1<br>**Reference AOP context**<br>**AOP 12:** Chronic binding of antagonist to N-methyl-D-aspartate receptors (NMDARs) during brain development leads to neurodegeneration with impairment in learning and memory in aging<br>**AOP 13:** Chronic binding of antagonist to N-methyl-D-aspartate receptors (NMDARs) during brain development induces impairment of learning and memory abilities<br>**Common sequence:** Antagonist binding to NMDARs → NMDAR inhibition → decreased calcium influx → reduced BDNF.<br>**AOP 12 branch:** Increased cell injury/death → neuroinflammation → neurodegeneration → learning and memory impairment.<br>**AOP 13 branches:** Aberrant dendritic morphology, reduced presynaptic glutamate release, or increased cell injury/death → decreased synaptogenesis → decreased neuronal network function → learning and memory impairment.<br>Branches continue from reduced BDNF. Reference event names are shortened; all four recorded paths are represented. |  |
| **GPT-4o — zero-shot**<br>**MIE:** NMDA receptor (NMDAR) inhibition<br>**Model reasoning**<br>1. The query molecule shows high similarity to references inhibiting the NMDA receptor, suggesting it likely inhibits NMDAR.<br>2. NMDA receptor inhibition (E1) leads to reduced synaptic plasticity (E2), a key event in neuronal function.<br>3. Reduced synaptic plasticity (E2) impairs neuronal communication (E3), affecting cognitive processes. | **Gemini 3.5 Flash — zero-shot**<br>**MIE:** NMDA receptor (NMDAR) inhibition<br>**Model reasoning**<br>1. The query molecule contains bis-guanidinium/alkyl-guanidinium features very similar to reference compounds 5, 6, 7, and 8 (similarities ranging from 0.60 to 0.71), which are highly active as NMDA receptor inhibitors.<br>2. Inhibition of the NMDA receptor (E1) reduces calcium influx (E2) into postsynaptic neurons.<br>3. This decrease in intracellular calcium levels leads to reduced BDNF release and expression (E3), which is crucial for synaptic plasticity. |

|  |  |  |  |  |  |
| --- | --- | --- | --- | --- | --- |
| 4. Impaired neuronal communication (E3) results in learning and memory impairment (E4), a recognized adverse outcome.<br>**Predicted AOs:** Learning and memory impairment (AO:341) |  |  | 4. Insufficient BDNF and calcium signaling impair long-term potentiation (LTP) (E4), a cellular mechanism of memory formation, ultimately leading to impaired learning and memory (E5, AO:341).<br>**Predicted AOs:** Learning and memory impairment (AO:341) |  |  |
| Explanation assessment and reference matching |  |  |  |  |  |
| GPT-4o — zero-shot |  |  | Gemini 3.5 Flash — zero-shot |  |  |
| Dimension | Naive-LLM judge | Rubric-based LLM judge | Dimension | Naive-LLM judge | Rubric-based LLM judge |
| Hallucination avoidance | 10-Jun | 10-Apr | Hallucination avoidance | 10-Aug | 10-Aug |
| Causal coherence | 10-Jul | 10-Aug | Causal coherence | 10-Sep | 10-Sep |
| Biological fidelity | 10-Jun | 10-Jul | Biological fidelity | 10-Aug | 10-Aug |
| Informativeness | 10-May | 10-May | Informativeness | 10-Jul | 10-Aug |
| Overall | 10-Jun | 10-Jun | Overall | 10-Aug | 8.25/10 |
| **Reference matching**<br>MIE: 1/1; AO: 1/1; KE: 0/10; KER: 0/14<br>**Reference alignment summary**<br>1. Identifies NMDAR inhibition and the reference AO through a broad synaptic-plasticity explanation.<br>2. Omits calcium and BDNF intermediates; neither structured KE matches under automated event mapping. |  |  | **Reference matching**<br>MIE: 1/1; AO: 1/1; KE: 2/10; KER: 2/14<br>**Reference alignment summary**<br>1. Specifies decreased calcium influx and reduced BDNF, recovering two reference KEs and two KERs.<br>2. Adds long-term potentiation, which is absent from the selected reference paths; pathway recovery remains incomplete. |  |  |

The upper block presents the reference AOP context, model reasoning, and predicted adverse outcomes. The lower block reports reference-matching results, scores from both LLM-judge configurations, and pathway-alignment summaries.

### 3.3. Few-shot prompting

Few-shot prompting improved AO prediction and downstream pathway matching in both GPT-5.1 and Gemini 3.8 Flash (Table 5). AO, KE, and KER performance increased across the one-, two-, and four-shot conditions, with the largest gains occurring between two and four examples. From zero to four shots, AO macro-F1 increased from 0.424 to 0.792 for GPT-5.1 and from 0.463 to 0.956 for Gemini 3.8 Flash. Under the four-shot condition, KE and KER macro-F1 reached 0.359 and 0.262 for GPT-5.1 and 0.492 and 0.457 for Gemini 3.8 Flash, respectively. Examples therefore increased agreement with both developmental outcomes and the downstream pathway annotations linking molecular perturbations to those outcomes. Nevertheless, KE and KER agreement remained lower than AO agreement, indicating that improved outcome prediction did not ensure comparable recovery of the annotated pathway structure.

Several factors may have contributed to these improvements. Each example included the chemical reference context, reference MIE and AO labels, structured AOP, and a completed mechanistic explanation. Beyond illustrating the task, the examples exposed the models to reference terminology, expected event specificity, output structure, and MIE-to-pathway mappings. Improved KE and KER performance may consequently reflect both better pathway inference and closer alignment with the terminology and structure used in the reference annotations.

The effects on MIE prediction differed between models. GPT-5.1 increased from a zero-shot MIE macro-F1 of 0.614 to 0.774 with four examples, although performance was not monotonic across the intermediate conditions. Gemini 3.8 Flash showed little change from its higher zero-shot baseline, increasing from 0.804 to 0.813. Thus, few-shot prompting improved both initiating-event and downstream reference agreement for GPT-5.1, whereas the gains for Gemini 3.8 Flash occurred predominantly in AO, KE, and KER matching.

Overall explanation scores also increased with examples under both LLM-judge configurations (Tables 6 and 7). From zero to four shots, scores under the naive-LLM judge increased from 4.479 to 6.845 for GPT-5.1 and from 5.123 to 7.012 for Gemini 3.8 Flash. Corresponding scores under the rubric-based LLM judge increased from 4.735 to 6.639 and from 5.187 to 6.911, respectively. For GPT-5.1, gains in causal coherence, biological fidelity, and hallucination avoidance exceeded the gain in informativeness under both configurations. Thus, examples improved several judged aspects of explanation quality alongside agreement with the reference annotations.

The progressive improvements observed here contrast with the few-shot pattern reported in ToxReason, where Qwen3-4B achieved its highest prediction and explanation performance with one example and declined with additional examples ^23^. Indeed, additional examples do not necessarily improve performance, and their effectiveness can depend on which examples are selected and the model receiving them ^43, 44^. In our study, each example combined chemical reference context, pathway annotations, and a completed mechanistic response, so the contribution of these components cannot be separated. Moreover, the one- and two-shot conditions were subsets of the four-example set.

Finally, we compared our few-shot LLM models with a supervised chemical structure-based baseline using ECFP and RF model. The ECFP4-RF achieved an AO macro-F1 of 0.915, compared with AO macro-F1 of 0.792 for GPT-5.1 and 0.956 for Gemini 3.8 Flash under four-shot prompting. These values arise from different learning settings: the random forest was trained directly on a large set of chemical AO labels, whereas the LLMs received in-context examples together with analogue target-activity evidence. An additional distinction is that the ECFP4-RF generated endpoint predictions only, while the LLMs also produced explicit MIE–KE–AO explanations that allowed the proposed biological reasoning behind each prediction to be examined.

### 3.4. Sensitivity of explanation assessment to LLM-judge configuration

The two LLM-judge configurations produced generally consistent rankings of explanations across compounds within each model and prompting condition. Overall, Spearman correlations ranged from 0.835 to 0.975 (Figure 2a). However, similar rankings did not imply identical scoring profiles. Across all 16 conditions, the rubric-based LLM judge assigned lower mean scores for hallucination avoidance and higher mean scores for causal coherence and informativeness than the Naive-LLM judge (Tables 6 and 7). Biological fidelity scores were higher in 14 conditions and lower in two. These opposing shifts partly offset one another in the overall score. Mean paired differences, calculated as rubric-based LLM judge minus Naive-LLM judge, ranged from −0.207 to 0.256 points (Figure 2b). Thus, similar overall scores could mask systematic differences in how individual dimensions were assessed. Both configurations used the same judge model, paired inputs, and permitted score range. The comparison therefore demonstrates sensitivity to rubric instructions but does not establish the accuracy of either judge.

**Figure 2.**
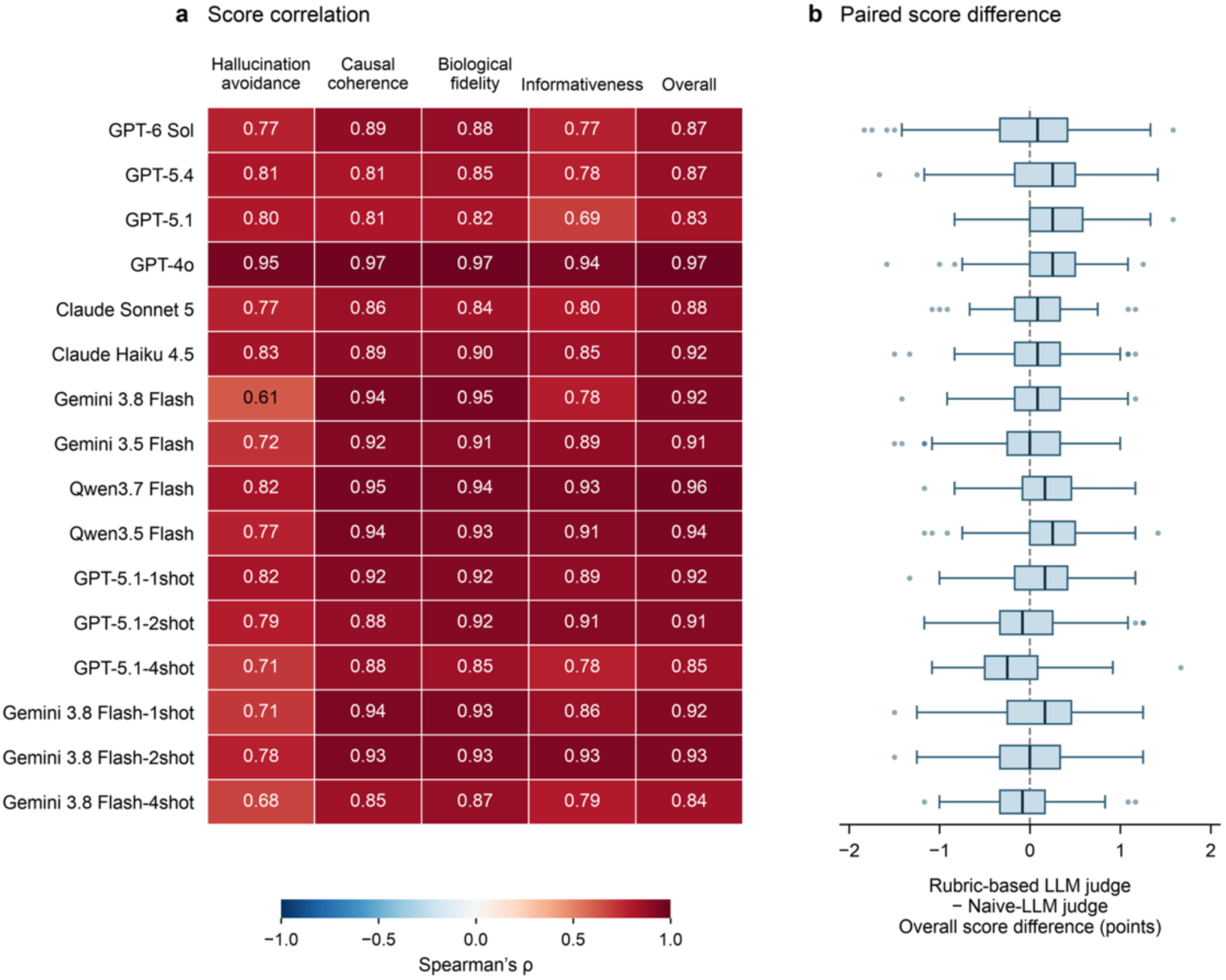
Comparison of the Naive-LLM judge and Rubric-based LLM judge. (a) Spearman correlations for four dimensions and their mean (Overall). (b) Paired Overall score differences (Rubric-based LLM judge − Naive-LLM judge). Scores were averaged across three repeats per chemical (n = 251 per model condition). Boxes show the median and interquartile range; whiskers extend to the most extreme values within 1.5 times the interquartile range, with remaining values shown as points.

### 3.5. Case study: comparison of LLM reasoning

We used two chemical-level case studies to illustrate complementary aspects of model performance across prompting conditions and models. The first case study compared zero- and four-shot responses from the same model to examine the effect of few-shot prompting, and the second compared zero-shot responses from different models to illustrate between-model differences in mechanistic explanation. For the chemical C^30^H^37^NO^4^ (CHEMBL3939687), Gemini 3.8 Flash identified retinoic acid receptor activation and both reference AOs under zero- and four-shot conditions. The zero-shot explanation mentioned impaired progenitor proliferation but described the downstream events more generally. With four examples, the response specified altered cell-cycle gene expression, reduced cortical neural progenitor proliferation, altered brain morphology, and reduced cortical thickness. Reference matching consequently increased from 0/4 KEs and 0/9 KERs to 4/4 KEs and 9/9 KERs, respectively. For the chemical C^16^H^28^Cl^2^N^6^O (CHEMBL117030), GPT-4o and Gemini 3.5 Flash both identified NMDAR inhibition and learning and memory impairment under zero-shot conditions. GPT-4o connected these through general changes in synaptic plasticity and neuronal communication, whereas Gemini 3.5 Flash additionally specified decreased calcium influx and reduced BDNF, matching two reference KEs and two KERs. The latter explanation also introduced long-term potentiation, which was not represented in the selected reference pathways, and downstream pathway recovery remained incomplete. These examples illustrate that agreement at the MIE and AO levels can coexist with substantial differences in the specificity and reference alignment of downstream mechanistic explanations.

The upper block presents the reference AOP context, model reasoning, and predicted adverse outcomes. The lower block reports reference-matching results, scores from both LLM-judge configurations, and pathway-alignment summaries.

### 3.6. Environmental Implications, Limitations, and Future Directions

Experimental DNT assessment is costly, time-consuming, and difficult to apply across large chemical inventories. AOP-informed LLMs offer a complementary approach for integrating chemical activity information with structured biological knowledge and generating explicit MIE-KE-AO hypotheses. LLMs can therefore be evaluated not only on which adverse outcomes they predict but also on how those outcomes may arise. This distinction will become increasingly important as artificial intelligence is incorporated into NAMs ^45, 46^. However, the generated explanations should be regarded as testable hypotheses rather than evidence of toxicity. Our analysis evaluates AOP-informed reasoning and does not establish compound-level DNT hazard. ChEMBL activity measurements and the 10 μM screening criterion do not account for exposure, bioavailability, or dose–response relationships, and the benchmark covers a limited set of AOs and AOPs. Pathway prediction scores also depend on matching generated descriptions to AOP terminology. With broader chemical and AOP coverage and experimental validation, AOP-informed LLMs could support more transparent and mechanism-informed DNT assessment as a complement to experimental testing.

## References

(1) Stiles, J.; Jernigan, T. L. The Basics of Brain Development. Neuropsychology Review 2010, 20 (4), 327–348. DOI: 10.1007/s11065-010-9148-4.

(2) Rice, D.; Barone, S., Jr. Critical periods of vulnerability for the developing nervous system: evidence from humans and animal models. Environ Health Perspect 2000, 108 *Suppl 3* (Suppl 3), 511–533. DOI: 10.1289/ehp.00108s3511 From NLM.

(3) Grandjean, P.; Landrigan, P. J. Neurobehavioural effects of developmental toxicity. The Lancet Neurology 2014, 13 (3), 330–338. DOI: 10.1016/S1474-4422(13)70278-3 (acccessed 2026/09/25).

(4) Fritsche, E.; Grandjean, P.; Crofton, K. M.; Aschner, M.; Goldberg, A.; Heinonen, T.; Hessel, E. V. S.; Hogberg, H. T.; Bennekou, S. H.; Lein, P. J.;, et al. Consensus statement on the need for innovation, transition and implementation of developmental neurotoxicity (DNT) testing for regulatory purposes. Toxicology and Applied Pharmacology 2018, 354, 3–6. DOI: 10.1016/j.taap.2018.02.004.

(5) Crofton, K. M.; Mundy, W. R.; Shafer, T. J. Developmental neurotoxicity testing: a path forward. Congenit Anom (Kyoto) 2012, 52 (3), 140–146. DOI: 10.1111/j.1741-4520.2012.00377.x From NLM.

(6) Mansouri, K.; Karmaus, A. L.; Fitzpatrick, J.; Patlewicz, G.; Pradeep, P.; Alberga, D.; Alepee, N.; Allen, T. E. H.; Allen, D.; Alves, V. M.;, et al. CATMoS: Collaborative Acute Toxicity Modeling Suite. Environmental Health Perspectives 2021, 129 (4). DOI: 10.1289/ehp8495 (acccessed 9/28/2026).

(7) Gao, F.; Zhang, W.; Baccarelli, A. A.; Shen, Y. Predicting chemical ecotoxicity by learning latent space chemical representations. Environ Int 2022, 163, 107224. DOI: 10.1016/j.envint.2022.107224 From NLM.

(8) Gao, F.; Wolf, G.; Hirn, M. Geometric Scattering for Graph Data Analysis. In Proceedings of the 36th International Conference on Machine Learning, Proceedings of Machine Learning Research; 2019.

(9) Huang, R.; Xia, M.; Sakamuru, S.; Zhao, J.; Shahane, S. A.; Attene-Ramos, M.; Zhao, T.; Austin, C. P.; Simeonov, A. Modelling the Tox21 10 K chemical profiles for in vivo toxicity prediction and mechanism characterization. Nature Communications 2016, 7 (1), 10425. DOI: 10.1038/ncomms10425.

(10) Sewell, F.; Alexander-White, C.; Brescia, S.; Currie, R. A.; Roberts, R.; Roper, C.; Vickers, C.; Westmoreland, C.; Kimber, I. New approach methodologies (NAMs): identifying and overcoming hurdles to accelerated adoption. Toxicol Res (Camb) 2024, 13 (2), tfae044. DOI: 10.1093/toxres/tfae044 From NLM.

(11) Mayr, A.; Klambauer, G.; Unterthiner, T.; Hochreiter, S. DeepTox: Toxicity Prediction using Deep Learning. Frontiers in Environmental Science 2016, *Volume* 3 *-* 2015, Original Research. DOI: 10.3389/fenvs.2015.00080.

(12) Jeong, J.; Choi, J. Artificial Intelligence-Based Toxicity Prediction of Environmental Chemicals: Future Directions for Chemical Management Applications. Environ Sci Technol 2022, 56 (12), 7532–7543. DOI: 10.1021/acs.est.1c07413 From NLM.

(13) Guo, W.; Liu, J.; Dong, F.; Song, M.; Li, Z.; Khan, M. K. H.; Patterson, T. A.; Hong, H. Review of machine learning and deep learning models for toxicity prediction. Exp Biol Med (Maywood) 2023, 248 (21), 1952–1973. DOI: 10.1177/15353702231209421 From NLM.

(14) Villeneuve, D. L.; Crump, D.; Garcia-Reyero, N.; Hecker, M.; Hutchinson, T. H.; LaLone, C. A.; Landesmann, B.; Lettieri, T.; Munn, S.; Nepelska, M.;, et al. Adverse outcome pathway (AOP) development I: strategies and principles. Toxicol Sci 2014, 142 (2), 312–320. DOI: 10.1093/toxsci/kfu199 From NLM.

(15) Bal-Price, A.; Crofton, K. M.; Sachana, M.; Shafer, T. J.; Behl, M.; Forsby, A.; Hargreaves, A.; Landesmann, B.; Lein, P. J.; Louisse, J.;, et al. Putative adverse outcome pathways relevant to neurotoxicity. Crit Rev Toxicol 2015, 45 (1), 83–91. DOI: 10.3109/10408444.2014.981331 From NLM.

(16) Villeneuve, D. L.; Crump, D.; Garcia-Reyero, N.; Hecker, M.; Hutchinson, T. H.; LaLone, C. A.; Landesmann, B.; Lettieri, T.; Munn, S.; Nepelska, M.;, et al. Adverse outcome pathway development II: best practices. Toxicol Sci 2014, 142 (2), 321–330. DOI: 10.1093/toxsci/kfu200 From NLM.

(17) Jablonka, K. M.; Schwaller, P.; Ortega-Guerrero, A.; Smit, B. Leveraging large language models for predictive chemistry. Nature Machine Intelligence 2024, 6 (2), 161–169. DOI: 10.1038/s42256-023-00788-1.

(18) Cai, X.; Lai, H.; Wang, X.; Wang, L.; Liu, W.; Wang, Y.; Wang, Z.; Cao, D.; Zeng, X. Comprehensive evaluation of molecule property prediction with ChatGPT. Methods 2024, 222, 133–141. DOI: 10.1016/j.ymeth.2024.01.004 From NLM.

(19) Zheng, Y.; Koh, H. Y.; Ju, J.; Nguyen, A. T. N.; May, L. T.; Webb, G. I.; Pan, S. Large language models for scientific discovery in molecular property prediction. Nature Machine Intelligence 2025, 7 (3), 437–447. DOI: 10.1038/s42256-025-00994-z.

(20) Xian, Z.; Gu, J.; Li, L.; Liang, S. MolRAG: Unlocking the Power of Large Language Models for Molecular Property Prediction. Vienna, Austria, July, 2025; Association for Computational Linguistics: pp 15513–15531. DOI: 10.18653/v1/2025.acl-long.755.

(21) Yang, H.; Xiu, J.; yan, W.; Liu, K.; Cui, H.; Wang, Z.; He, Q.; Gao, Y.; Han, W. Large Language Models as Tools for Molecular Toxicity Prediction: AI Insights into Cardiotoxicity. Journal of Chemical Information and Modeling 2025, 65 (5), 2268– 2282. DOI: 10.1021/acs.jcim.4c01371 (acccessed 9/28/2026).

(22) Park, J.; Park, Y.; Song, M.; Park, S.; Lee, D.; Baek, S.; Kang, J. CoTox: Chain-of-Thought-Based Molecular Toxicity Reasoning and Prediction. In 2025 IEEE International Conference on Bioinformatics and Biomedicine (BIBM), 15–18 Dec. 2025, 2025; pp 4002–4007. DOI: 10.1109/BIBM66473.2025.11356816.

(23) Park, J.; Jang, W.; Kim, C.; Park, Y.; Kang, J. ToxReason: A Benchmark for Mechanistic Chemical Toxicity Reasoning via Adverse Outcome Pathway. San Diego, California, United States, July, 2026; Association for Computational Linguistics: pp 19542–19564. DOI: 10.18653/v1/2026.findings-acl.977.

(24) Zdrazil, B.; Felix, E.; Hunter, F.; Manners, E. J.; Blackshaw, J.; Corbett, S.; de Veij, M.; Ioannidis, H.; Lopez, D. M.; Mosquera, Juan F.; et al. The ChEMBL Database in 2023: a drug discovery platform spanning multiple bioactivity data types and time periods. Nucleic Acids Research 2024, 52 (D1), D1180–D1192. DOI: 10.1093/nar/gkad1004 (acccessed 9/26/2026).

(25) Society for Advancement of AOPs. AOP-Wiki. 2026. (acccessed 2026/09/25).

(26) Gadaleta, D.; Garcia de Lomana, M.; Serrano-Candelas, E.; Ortega-Vallbona, R.; Gozalbes, R.; Roncaglioni, A.; Benfenati, E. Quantitative structure–activity relationships of chemical bioactivity toward proteins associated with molecular initiating events of organ-specific toxicity. Journal of Cheminformatics 2024, 16 (1), 122. DOI: 10.1186/s13321-024-00917-x.

(27) rdkit/rdkit: 2026_03_6 (Q1 2026) Release; Zenodo: 2026. 10.5281/zenodo.22140358 (accessed.

(28) Rogers, D.; Hahn, M. Extended-Connectivity Fingerprints. Journal of Chemical Information and Modeling 2010, 50 (5), 742–754. DOI: 10.1021/ci100050t (acccessed 9/26/2026).

(29) OpenAI. GPT-6 Sol. 2026. (acccessed 2026/09/25).

(30) OpenAI. Introducing GPT-5.4. 2026. (acccessed 2026/09/25).

(31) OpenAI. GPT-5.1 in ChatGPT. 2025. (acccessed 2026/09/25).

(32) OpenAI. Hello GPT-4o. 2024. (acccessed 2026/09/25).

(33) Anthropic. Introducing Claude Sonnet 5. 2026. (acccessed 2026/09/25).

(34) Anthropic. Introducing Claude Haiku 4.5. 2025. (acccessed 2026/09/25).

(35) Google, D. Gemini 3.8 Flash. 2026. (acccessed 2026/09/25).

(36) Google, D. Gemini 3.5 Flash. 2026. (acccessed 2026/09/25).

(37) Alibaba, C. qwen3.7-flash. 2026. (acccessed 2026/09/25).

(38) Alibaba, C. qwen3.5-flash. 2026. (acccessed 2026/09/25).

(39) Breiman, L. Random Forests. Machine Learning 2001, 45 (1), 5–32. DOI: 10.1023/A:1010933404324.

(40) Brown, T. B.; Mann, B.; Ryder, N.; Subbiah, M.; Kaplan, J. D.; Dhariwal, P.; Neelakantan, A.; Shyam, P.; Sastry, G.; Askell, A.;, et al. Language Models are Few-Shot Learners. In Advances in Neural Information Processing Systems, 2020.

(41) OpenAI. GPT-6 Astra: A new generation of intelligence. 2026. (acccessed 2026/09/25).

(42) OpenAI. Introducing GPT-5.4 mini and nano. 2026. (acccessed 2026/09/25).

(43) Chen, J.; Chen, L.; Zhu, C.; Zhou, T. How Many Demonstrations Do You Need for In-context Learning? Singapore, December, 2023; Association for Computational Linguistics: pp 11149–11159. DOI: 10.18653/v1/2023.findings-emnlp.745.

(44) Peng, K.; Ding, L.; Yuan, Y.; Liu, X.; Zhang, M.; Ouyang, Y.; Tao, D. Revisiting Demonstration Selection Strategies in In-Context Learning. Bangkok, Thailand, August, 2024; Association for Computational Linguistics: pp 9090–9101. DOI: 10.18653/v1/2024.acl-long.492.

(45) Schmeisser, S.; Miccoli, A.; von Bergen, M.; Berggren, E.; Braeuning, A.; Busch, W.; Desaintes, C.; Gourmelon, A.; Grafström, R.; Harrill, J.;, et al. New approach methodologies in human regulatory toxicology - Not if, but how and when! Environ Int 2023, 178, 108082. DOI: 10.1016/j.envint.2023.108082 From NLM.

(46) Kleinstreuer, N.; Hartung, T. Artificial intelligence (AI)—it’s the end of the tox as we know it (and I feel fine)*. Archives of Toxicology 2024, 98 (3), 735–754. DOI: 10.1007/s00204-023-03666-2.

